# Widespread destabilization of *C. elegans* microRNAs by the E3 ubiquitin ligase EBAX-1

**DOI:** 10.1101/2024.06.28.601170

**Authors:** Michael W. Stubna, Aditi Shukla, David P. Bartel

## Abstract

MicroRNAs (miRNAs) associate with Argonaute (AGO) proteins to form complexes that direct mRNA repression. miRNAs are also the subject of regulation. For example, some miRNAs are destabilized through a pathway in which pairing to specialized transcripts recruits the ZSWIM8 E3 ubiquitin ligase, which polyubiquitinates AGO, leading to its degradation and exposure of the miRNA to cellular nucleases. Here, we found that 22 miRNAs in *C. elegans* are sensitive to loss of EBAX-1, the ZSWIM8 ortholog in nematodes, implying that these 22 miRNAs might be subject to this pathway of target-directed miRNA degradation (TDMD). The impact of EBAX-1 depended on the developmental stage, with the greatest effect on the miRNA pool (14.5%) observed in L1 larvae and the greatest number of different miRNAs affected (17) observed in germline-depleted adults. The affected miRNAs included the miR-35–42 family, as well as other miRNAs among the least stable in the worm, suggesting that TDMD is a major miRNA-destabilization pathway in the worm. The excess miR-35–42 molecules that accumulated in *ebax-1* mutants caused increased repression of their predicted target mRNAs and underwent 3′ trimming over time. In general, however, miRNAs sensitive to EBAX-1 loss had no consistent pattern of either trimming or tailing. Replacement of the 3′ region of miR-43 substantially reduced EBAX-1 sensitivity, a result that differed from that observed previously for miR-35. Together, these findings broaden the implied biological scope of TDMD-like regulation of miRNA stability in animals, and indicate that a role for miRNA 3′ sequences is variable in the worm.

## INTRODUCTION

MicroRNAs (miRNAs) are ∼22-nt RNAs that associate with Argonaute (AGO) proteins to direct the posttranscriptional repression of mRNAs (Jonas and Izaurralde 2015; Bartel 2018). Within the miRNA–AGO complex, the miRNA recognizes sites within mRNAs, typically through pairing to the miRNA seed region (miRNA nucleotides 2–8) (Bartel 2009), and AGO recruits mRNA-deadenylation machinery, ultimately leading to destabilization and/or translational repression of the targeted mRNA (Jonas and Izaurralde 2015). More than 500 miRNAs have been identified in humans (Fromm et al. 2015). Together, they regulate most human mRNAs (Friedman et al. 2009), and many of the broadly conserved mammalian miRNAs are essential for viability or proper development (Bartel 2018).

Most miRNAs are long-lived within the cell, with half-lives often exceeding a day (Kingston and Bartel 2019; Reichholf et al. 2019), presumably because their association with an AGO protein shields them from cellular nucleases. However, some miRNAs turn over more rapidly, with half-lives of only a few hours (Rissland et al. 2011; Kingston and Bartel 2019; Reichholf et al. 2019). Some of these relatively short-lived miRNAs are destabilized through the action of unusual target sites that direct their degradation. In this target-directed miRNA degradation (TDMD), pairing to the site recruits the ZSWIM8 Cullin/RING-type E3 ubiquitin ligase, leading to polyubiquitination and degradation of AGO, leaving the miRNA exposed to cytoplasmic nucleases (Han et al. 2020; Shi et al. 2020; Han and Mendell 2023; Buhagiar and Kleaveland 2024).

TDMD was first described as a phenomenon triggered by sites within viral or synthetic target RNAs (Ameres et al. 2010; Buck et al. 2010; Cazalla et al. 2010; Libri et al. 2012; de la Mata et al. 2015). More recently, cellular transcripts have also been found to have sites that direct miRNA degradation. For example, a site in the murine *Nrep* 3′ UTR—conserved to the long noncoding (lnc)RNA *libra* in zebrafish—directs degradation of miR-29b (Bitetti et al. 2018), and a site in the Cyrano lncRNA triggers destruction of miR-7 (Kleaveland et al. 2018). Two other sites that direct miRNA degradation have been reported among mammalian transcripts (Ghini et al. 2018; Simeone et al. 2022), and another six have been found among fly transcripts (Kingston et al. 2022; Sheng et al. 2023). Each of these 10 sites reported within endogenous TDMD triggers resembles sites reported within viral or artificial triggers in having not only pairing to the miRNA seed but also extensive pairing to the miRNA 3′ region. This pairing to the miRNA 3′ region causes conformational changes to the protein and mRNA (Sheu-Gruttadauria et al. 2019), which are proposed to favor binding of ZSWIM8 and polyubiquitination of AGO (Han et al. 2020; Shi et al. 2020).

Many additional cases of endogenous TDMD are also thought to occur, as indicated by the 30 named miRNAs that increase upon loss of ZSWIM8 in mammalian cells (Shi et al. 2020), the 74 miRNAs reported to increase upon ZSWIM8 loss in diverse mouse tissues (Jones et al. 2023; Shi et al. 2023), the 21 miRNAs that increase upon loss of the ZSWIM8 ortholog in Drosophila S2 cells or embryos (Shi et al. 2020; Kingston et al. 2022), and the 10 miRNAs that increase upon loss of the ZSWIM8 ortholog in *Caenorhabditis elegans* gravid adults (Shi et al. 2020). Indeed, the magnitude of increase observed upon ZSWIM8 loss corresponds closely with miRNA half-life, implying that TDMD explains the short half-lives of most short-lived miRNAs in mammalian and Drosophila cells (Shi et al. 2020).

In *C. elegans* gravid adults, the miRNAs that increase upon loss of the ZSWIM8 ortholog, known as EBAX-1, include miRNAs of the miR-35 family (Shi et al. 2020). The eight members of this family are highly expressed in embryos and required for embryonic development (Alvarez-Saavedra and Horvitz 2010; Dexheimer et al. 2020). The observation that members of the miR-35 family are EBAX-1 sensitive and rapidly decay as development proceeds to the first larval stage, implies that they are endogenous TDMD substrates (Shi et al. 2020; Donnelly et al. 2022). However, a *C. elegans* transcript that directs degradation of these miRNAs (or of any of the other EBAX-1–sensitive miRNAs) has yet to be reported. Moreover, the machinery that degrades miR-35 appears to recognize the miRNA seed but not its 3′ region, suggesting that if TDMD is occurring, it is specified through a process that, in stark contrast to all previously characterized examples, does not involve pairing to the miRNA 3′ end (Donnelly et al. 2022).

We set out to more fully understand the potential scope of endogenous TDMD in *C. elegans*, extending our analyses beyond gravid adults to identify which miRNAs are sensitive to EBAX-1 loss at each stage of worm development. We also examined whether the unusual 3′-region-independent recognition observed for miR-35 generalizes to other EBAX-1–sensitive miRNAs in *C. elegans*.

## RESULTS

### EBAX-1 appears to destabilize 22 miRNAs over the course of development

Because many *C. elegans* miRNAs are under temporal control (Lim et al. 2003; Kato et al. 2009), we reasoned that the EBAX-1–sensitive miRNAs reported in gravid adults (Shi et al. 2020) might be only a subset of those that are EBAX-1 sensitive over the course of development. To this end, we used small-RNA sequencing (sRNA-seq) to quantify the changes in miRNAs and other small RNAs upon EBAX-1 loss, comparing results for *ebax-1(tm2321)* null-mutant animals to those of WT animals across early and late embryos, four larval stages, and *glp-4(ts)* adults (which have greatly diminished germline). For comparison, we also reanalyzed the published datasets from gravid adults (Shi et al. 2020). To designate miRNAs as EBAX-1 sensitive, we combined DESeq2 differential expression analysis (Love et al. 2014) with a statistical model of experimental noise and secondary effects, which was based on the expectation that TDMD loss increases the levels of TDMD substrates, whereas the accompanying noise and secondary effects both decrease and increase miRNA levels (Wang and Bartel 2022).

At each developmental stage examined, at least one miRNA accumulated in *ebax-1* mutants (Fig. 1A). In sum, 22 unique miRNAs from nine miRNA families were EBAX-1 sensitive in at least one stage (Fig. 1B). L1 larvae homozygous for an independent null allele of *ebax-1*, *ju699*, had a pattern of miRNA accumulation closely matching that of *ebax-1(tm2321)* L1 larvae (Fig. S1D, S1A), which confirmed both the role of *ebax-1* in influencing miRNA levels and the reproducibility of our sRNA-seq analyses.

**Fig. 1.**
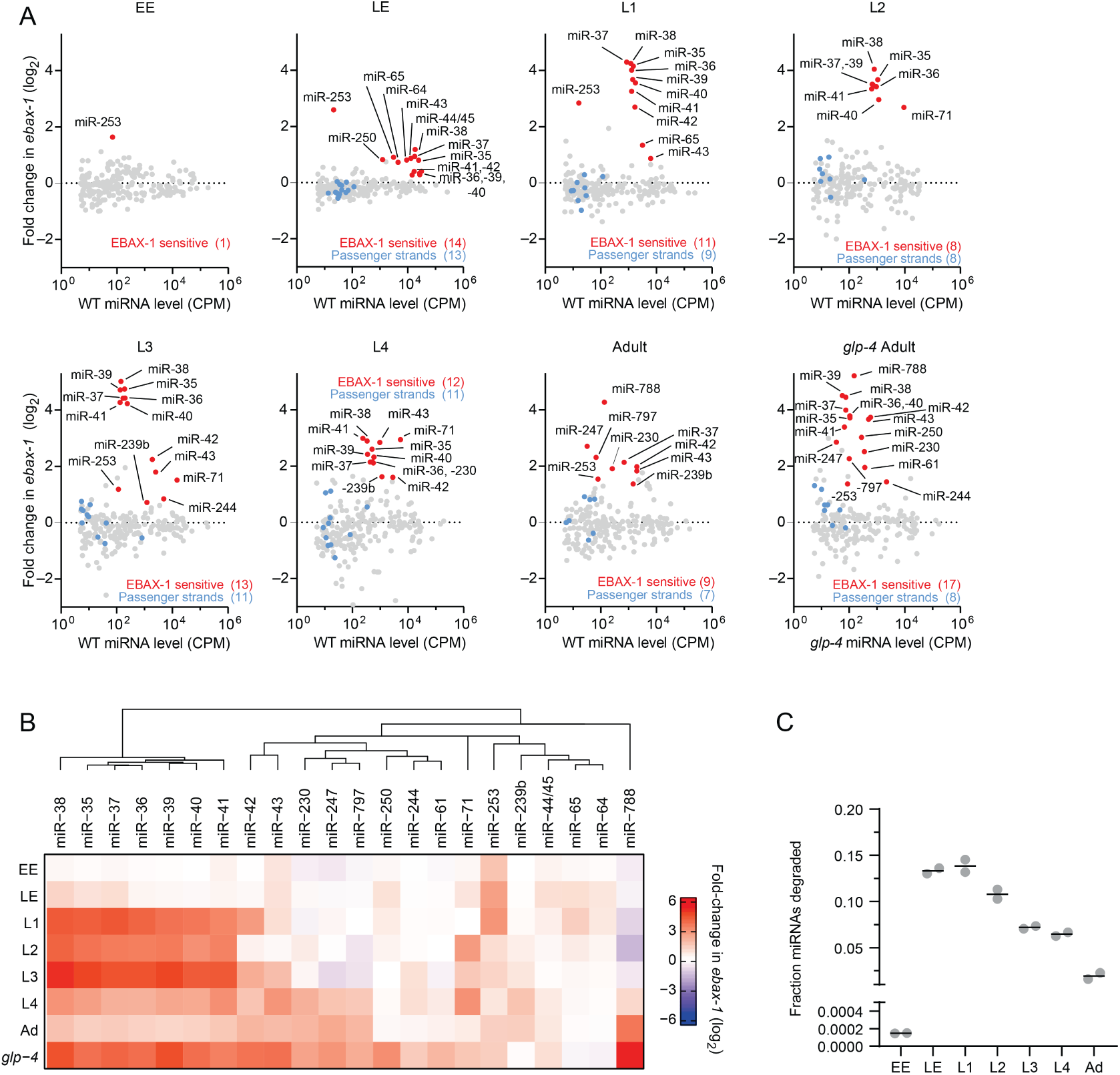
EBAX-1 limits accumulation of 22 miRNAs across worm development. **(A)** The effect of EBAX-1 loss on miRNA levels in worms of the indicated stage, as measured by sRNA-seq. Plotted are fold-changes in miRNA levels in *ebax-1(tm2321)* null-mutant worms compared to wild-type worms. Points for miRNAs called as EBAX-1–sensitive are red, and those of their corresponding passenger strands are blue. miRNAs were called as EBAX-sensitive by a statistical pipeline combining DESeq2 and a bi-beta uniform mixture (BBUM) model of noise and secondary effects with a false discovery rate (FDR) threshold of < 0.01 (Table S1). miRNA levels are expressed as counts per million miRNA reads (CPM; including one pseudocount) averaged across two biological replicates. miRNAs are filtered for a mean wild-type expression of ≥ 5 CPM (including the pseudocount) for reliable quantification. Some miRNAs did not have passenger strands that exceeded this expression cutoff, whereas miR-44/45 had two quantifiable passenger strands at each stage examined (Table S1). Results from adult worms are reanalyzed from Shi et al. (2020). For consistency, we reanalyzed only the two adult replicates whose *ebax-1* mutant samples are derived from strain CZ9907, as samples of all other stages were obtained using this strain. Our lower limit for the *y* axes excluded three points: miR-1817 in L4, miR-77 in L2, and lin-4 in EE. **(B)** Summary of EBAX-1–sensitive miRNAs identified through sRNA-seq. Shown are mean fold changes in miRNA levels upon EBAX-1 loss at each stage examined for miRNAs that are EBAX-sensitive at one or more stages. To group miRNAs with similar patterns of EBAX-1 sensitivity across development, we performed unsupervised hierarchical clustering on the fold-change data using the R package ComplexHeatmap with default clustering parameters. **(C)** The fraction of cellular miRNA molecules degraded through EBAX-1 across development. For each sRNA-seq library at each developmental stage, raw miRNA counts corresponding to EBAX-1–sensitive or non-EBAX-1–sensitive miRNAs were summed. Wild-type miRNA counts were then scaled so that the sum of their non-EBAX-1–sensitive miRNA counts equaled that in the *ebax-1* mutant in the same replicate, allowing absolute comparisons of EBAX-1–sensitive miRNA counts. The wild-type sum of EBAX-1–sensitive miRNA counts after scaling was then subtracted from the sum of unscaled EBAX-1–sensitive miRNA counts, yielding an estimate of how many EBAX-1–sensitive molecules accumulated in the *ebax-1* mutant. The resulting difference was divided by the unscaled sum of all miRNA counts in the *ebax-1* mutant library to yield the fraction of cellular miRNA molecules degraded. Each point denotes a biological replicate, and lines denote the mean of the two replicates.

To speak to whether this enhanced accumulation in the *ebax-1* strains represented impaired miRNA degradation or increased miRNA production, we examined the levels of miRNA passenger strands. miRNA passenger strands are produced in parallel with the miRNA strands as a side product of miRNA biogenesis (Bartel 2018). For the 22 miRNAs that significantly increased upon EBAX-1 loss at one or more stages, we observed relatively small changes in passenger-strand abundance (Fig. 1A, Fig. S1B), which indicated that miRNA production was not substantially altered—consistent with a post-transcriptional effect of EBAX-1 on mature, AGO-loaded miRNAs. Thus, our analyses of worms over the course of development increased the number of known EBAX-1–sensitive miRNAs from 10 to 22, with a concomitant increase in the implied scope of TDMD in nematodes.

Our results from L3 and L4 larvae were in substantial agreement with a recent report of miRNAs that increase upon EBAX-1 loss at mid-to-late larval stages (Nahar et al. 2024). Any differences with the other study were difficult to evaluate because it was carried out for other purposes, after culture at an elevated temperature (25°C), and did not report the fate of passenger strands or perform biological replicates required for statistical analysis.

*ebax-1* mRNA is expressed throughout development, with a modest peak during late embryogenesis (Fig. S1C) (Wang et al. 2013; Boeck et al. 2016). Fourteen miRNAs were sensitive to loss of *ebax-1* in late embryos (Fig. 1A, B). These included miR-43, miR-253, and miRNAs of the miR-35 family (miR-35–42), which had been identified as EBAX-1 sensitive in gravid adults (Shi et al. 2020), as well as miR-44/45, miR-64, miR-65, and miR-250, which were newly identified as EBAX-1 sensitive (Fig. 1A, B). All but three of these miRNAs continued to be classified as EBAX-1 sensitive into the L1 larval stage or beyond, with the change between mutant and wild-type for some miRNAs continuing to increase as worms developed. For example, miR-35 was elevated ∼1.7-fold in *ebax-1* late embryos and 18-fold in L1 larvae (Fig. 1A, B).

Later in larval development, additional miRNAs emerged as EBAX-1 sensitive. For example, miR-239b, which rapidly accumulates following heat shock (Schreiner et al. 2019), was elevated in mutant L3 larvae, and miR-230, another heat-responsive miRNA, was elevated at the L4 stage, and three other heat-responsive miRNAs, miR-247, miR-788, and miR-797 (Schreiner et al. 2019), were elevated in adults (Fig. 1A) (Shi et al. 2020).

Because gravid adults contain a substantial number of germline cells and embryos, which might obscure the role of EBAX-1 in the adult soma, our analyses included germline-depleted adults. When grown at a restrictive temperature of 25°C, worms homozygous for the *glp-4(bn2)* allele generate only 12 of the ∼1200 germ nuclei present in wild-type adults (Beanan and Strome 1992). Accordingly, we performed sRNA-seq on *glp-4(bn2);ebax-1(tm2321)* adults raised at the restrictive temperature and compared these results to those of the parental *glp-4(bn2)* strain cultured in parallel. Confirming the near-complete loss of germline, germline-enriched small RNAs, including miR-35–42, miR-43, and miR-70 (McEwen et al. 2016) and Piwi-interacting RNAs (piRNAs/21U-RNAs) (Ruby et al. 2006; Batista et al. 2008) were strongly depleted in samples from both the *glp-4;ebax-1* and the *glp-4* worms (Fig. S2A, B).

Seventeen miRNAs were sensitive to EBAX-1 loss in the *glp-4* background. These included all but one of the nine EBAX-1–sensitive miRNAs identified in germline-intact adults, as well as remaining members of the miR-35 family (miR-35, -36, -38, -39, -40, -41), miR-244, miR-250, and miR-61 (Fig. 1A, 1B). For each of these miRNAs identified in germline-depleted adults but not in germline-intact adults, levels increased in *ebax-1* germline-intact adults but did not satisfy our cutoff for statistical significance. Moreover, the fold upregulation of these miRNAs in germline-deficient adults was consistently greater than that in germline-intact adults (Fig. S2C), presumably because miRNA molecules accumulating in *glp-4(bn2)* animals were not diluted by newly transcribed molecules produced in germline and embryos. For example, loss of EBAX-1 elevated miR-61 ∼1.5-fold in germline-intact adults and ∼4-fold in germline-deficient adults (Fig. S2C). Importantly, most of these EBAX-1–sensitive miRNAs were among the least stable miRNAs of adult worm(Lehrbach et al. 2012) (Fig. S2D), implying that the action of EBAX-1 underlies the instability of most short-lived miRNAs of worms, as observed in mammalian and fly cells (Shi et al. 2020).

Having identified numerous EBAX-1–sensitive miRNAs, including some that were highly expressed, we wondered about the fraction of the cellular miRNA pool that is turned over through EBAX-1. At each developmental stage, we estimated the fraction of cellular miRNAs degraded due to EBAX-1 (Fig. 1C), using a normalization strategy that equalized the sum of non-EBAX-1–sensitive miRNA counts observed between wild-type and *ebax-1* mutant samples, enabling estimates of surplus miRNA molecules that accumulated in *ebax-1* mutants. In early embryos, miR-253, the only EBAX-1–sensitive miRNA, is expressed at a relatively low level (Fig. 1A), and thus its EBAX-1–dependent turnover affected less than 0.02% of embryonic miRNA molecules (Fig. 1C). By contrast, as much as 13.6% and 14.5% of miRNA molecules appeared to have turned over through EBAX-1 in late embryo and L1 larva, respectively, reflecting the abundance and strong EBAX-1 sensitivity of the miR-35 family (Fig. 1C). As the abundance of the miR-35 family miRNAs decreased over the course of larval development, so too did the proportion of cellular miRNAs made up of miR-35 family members, which resulted in progressively lower fractions of miRNAs degraded (Fig. 1C). These results indicated that, in terms of the fraction of miRNA molecules, the major wave of EBAX-1–dependent miRNA degradation occurs early in development, in line with the temporal expression pattern of *ebax-1* mRNA (Fig. S1C).

### EBAX-1 has little effect on piRNAs

Looking beyond miRNAs, we also examined how piRNAs responded to loss of EBAX-1 in adult worms. piRNAs associate with a PIWI-related Argonaute protein, PRG-1, in the worm germline to direct transposon silencing and promote gametogenesis (Weick and Miska 2014; Youngman and Claycomb 2014). Worm piRNAs tend to pair to their targets throughout the piRNA length, often with one or more mismatches in the central region (Shen et al. 2018; Zhang et al. 2018)— a pairing configuration sometimes associated with TDMD in mammals (Sheu-Gruttadauria et al. 2019) and flies (Ameres et al. 2010). However, no piRNAs were significantly upregulated in *ebax-1* adults (Fig. S3A, Table S2). We speculate that piRNAs are refractory to TDMD because the PIWI-related protein is not susceptible to EBAX-1–mediated polyubiquitination, as proposed for the Drosophila AGO paralog that associates with small interfering RNAs (Kingston and Bartel 2021).

Curiously, instead of upregulated piRNAs, we found a cohort of 57 downregulated piRNAs that cleared an adjusted *p*-value threshold of < 0.05, as well as hundreds more that failed to clear this threshold but nonetheless appeared downregulated by 2–6-fold (Fig. S3A, Table S2). Almost all of these downregulated piRNAs originated from a single 0.46 Mb region on chromosome IV, a subdomain of a large, dense cluster of piRNA genes (Fig. S3B, S3C). The observation that the downregulated piRNAs mapped predominantly to this single region of the genome strongly suggested that downregulation occurred at the transcriptional rather than the post-transcriptional level. Interestingly, expression of 46 protein-coding genes annotated within this region was unchanged by loss of *ebax-1* (Fig. S3D, Table S3), indicating a piRNA-specific transcriptional defect rather than a more general epigenetic anomaly in this region of the *ebax1(tm2321)* genome.

We suspected that this piRNA anomaly was unique to the *ebax-1(tm2321)* line and independent of *ebax-1* function. To test this idea, we examined piRNA abundance in *ebax-1(ju699)* gravid adults. The piRNAs downregulated in the *ebax-1(tm2321)* line, originating from genomic interval ChrIV:4,760,000–5,220,000, were not similarly downregulated in the *ebax-1(ju699)* line (Fig. S3F). These findings pointed to an *ebax-1–*independent, second-site mutation causing reduced transcription of a set of genomically adjacent piRNAs in the *ebax-1(tm2321)* line.

### *ebax-1* is required to clear miR-35–42 from the developed embryo

The EBAX-1–sensitive miRNAs included all eight members of the miR-35 family (Fig. 1A), consisting of miR-35–42 (Fig. 2A)(Lau et al. 2001). These miRNAs are redundantly required for viability and embryonic patterning (Alvarez-Saavedra and Horvitz 2010; Dexheimer et al. 2020) and promote multiple aspects of postembryonic development (Liu et al. 2011; Massirer et al. 2012; McJunkin and Ambros 2014; McJunkin and Ambros 2017; Benner et al. 2019; Tran et al. 2019). Seven of the miR-35 family members (miR-35–41) are processed from the same primary transcript, whereas miR-42 originates from a different transcript, enabling differential transcriptional control (Martinez et al. 2008; Alvarez-Saavedra and Horvitz 2010). Nonetheless, levels of all eight miRNAs of the family are high in early embryo and then diminish sharply at the embryo-to-L1 transition (Kato et al. 2009; Stoeckius et al. 2009; Donnelly et al. 2022).

**Fig. 2.**
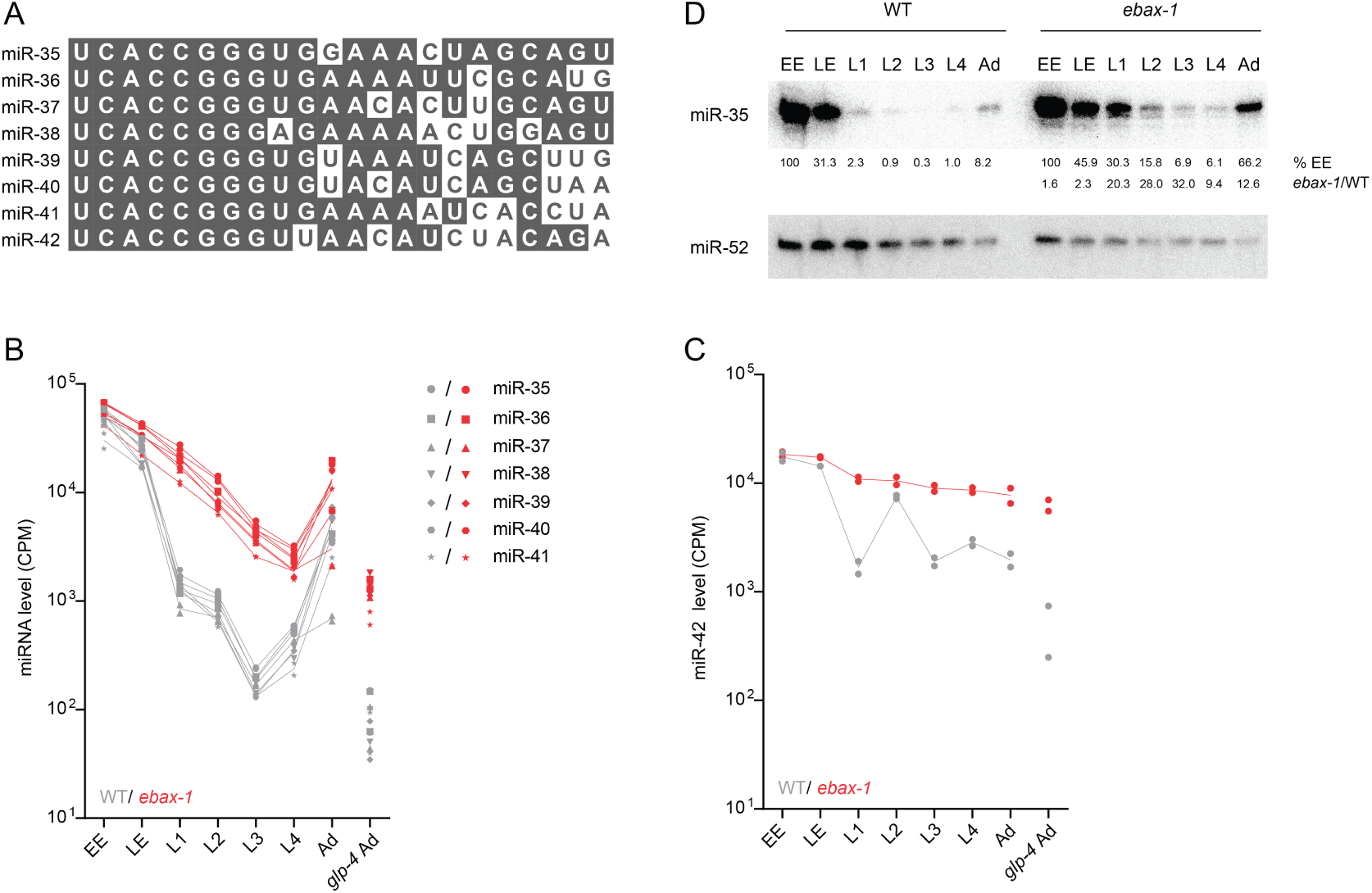
EBAX-1 is required to clear miR-35–42 at the embryo-to-L1 transition. **(A)** Sequence alignment of the eight members of the miR-35 family. **(B, C)** The effect of EBAX-1 on miR-35–42 levels across development. Expression of miR-35–42 family members in wild-type worms at each stage of development is shown in gray. Expression in *ebax-1(tm2321)* worms is plotted in red. Values for each of two replicates are plotted as separate points, whereas lines connect means of replicates. Expression of miR-42 is plotted separately as this miRNA is produced from a different primary transcript than that of miR-35–41. **(D)** The influence of EBAX-1 on miR-35 levels. Shown is an RNA blot comparing miR-35 and miR-52 levels in wild-type and *ebax-1* mutant worms at the stages indicated. Levels of miR-35 were normalized to those of miR-52 and then to levels of EE (%EE). Levels in *ebax-1* were also normalized to those of WT (*ebax-1*/WT).

To determine the consequences of EBAX-1 loss on miR-35–42 abundance at the embryo-to-L1 transition, we examined our sRNA-seq datasets (Table S1). As expected, wild-type miR-35–42 levels peaked in early embryo and declined precipitously at the embryo-to-L1 transition (Fig. 2B, 2C). Loss of EBAX-1 eliminated the precipitous drop of these eight miRNAs at this transition (Fig. 2B, 2C), supporting the proposal that EBAX-1 is required for the rapid clearing of these embryonic miRNAs that normally occurs at this time (Shi et al. 2020).

These results extended those of previous studies, which observe accumulation of some miR-35–42 miRNAs in either *ebax-1* gravid adults or *ebax-1* L1 larvae (Shi et al. 2020; Donnelly et al. 2022), a difference being that our current analysis found larger, more reproducible differences that achieved statistical significance for all eight family members. Indeed, the uniform effects observed at the L1 stage for all eight family members (Fig. 2B, 2C), despite their divergent 3′ regions (Fig. 2A), bolstered the conclusion that EBAX-dependent targeted degradation of this family occurs through an unusual mechanism that does not require recognition of the miRNA 3′ region (Donnelly et al. 2022).

After the first larval stage, the behavior of the miR-35–41 and miR-42 miRNAs diverged (Fig. 2B, 2C). Over the course of larval development, levels of miR-35–41 in wild-type worms and in *ebax-1* worms fell roughly in parallel with each other until stage L4, when production of these miRNAs resumed (Fig. 2B). This renewed production was presumably in the germline, as decline in residual miR-35–41 continued unabated in the *glp-4(bn2)* background at restrictive temperature (Fig. 2B), a context that substantially reduces production of miR-35 family members (Lau et al. 2001; McEwen et al. 2016). In contrast, the difference in miR-42 levels observed between wild-type and *ebax-1* worms narrowed at L2, presumably because miR-42 production transiently increased at this earlier stage, which muted the decline of this miRNA over the course of development in both wild-type and *ebax-1* animals (Fig. 2C).

Northern-blot analysis corroborated these findings, showing, for example, that loss of EBAX-1 in L1 larvae caused a ∼20-fold increase in miR-35 levels (Fig. 2D). Also in line with our sRNA-seq results, miR-35 persisted at higher levels in the mutant than in the wild-type throughout larval development. For example, in mutant L2 larvae, miR-35 was present at ∼16% of its early-embryonic level, whereas in wild-type L2 larvae it dropped to ∼0.9% of its early-embryonic level (Fig. 2D). These findings support the proposal that EBAX-1 is responsible for the rapid destabilization of miR-35–42 at the embryo-to-L1 transition (Shi et al. 2020; Donnelly et al. 2022).

### Accumulated miRNAs in *ebax-1* mutants dysregulate predicted mRNA targets

To investigate the possibility that miRNA accumulation observed in *ebax-1* mutants might inappropriately repress target mRNAs, we used poly(A)+ RNA-seq to quantify changes in mRNAs upon loss of EBAX-1 in L1 larvae. At an adjusted *p*-value threshold of < 0.05, this analysis identified 765 mRNAs that were downregulated and 289 that were upregulated (Fig. 3A, Table S4). Moreover, predicted targets of the miR-35 family tended to be repressed in the in *ebax-1* mutant larvae (Fig. 3B). As expected for authentic regulatory interactions (Friedman et al. 2009), repression was greater for predicted targets with sites classified as conserved among nematodes (Jan et al. 2011) (Fig. 3C). These observations indicate that the miR-35–42 molecules that persist in *ebax-1* larvae are functional and inappropriately continue to repress their targets after embryogenesis.

**Fig. 3.**
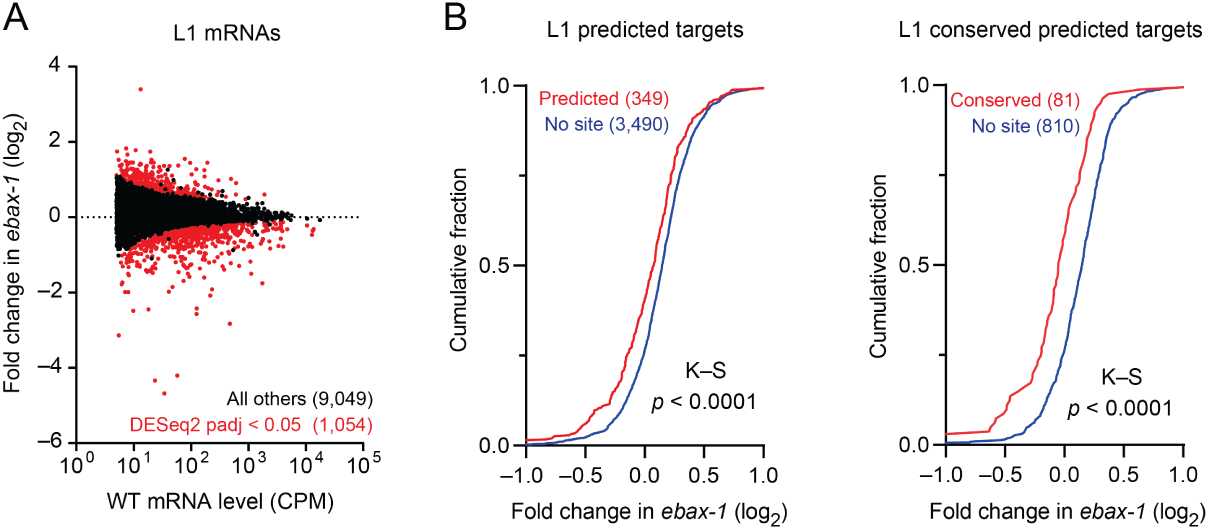
Loss of EBAX-1 causes increased repression of miR-35–42 target mRNAs. **(A)** The effect of EBAX-1 on mRNA levels in L1 larvae. Plotted are fold changes in mean mRNA levels (from two biological replicates) observed upon loss of EBAX-1 in L1 larvae. The mRNA levels are expressed as counts per million mRNA reads (CPM) and filtered for those with mean wild-type expression level ≥ 5 CPM. Points for differentially regulated mRNAs meeting a DESeq2 adjusted *p-*value threshold of < 0.05 are colored in red. **(B)** The influence of EBAX-1 on the expression of predicted mRNA targets of miR-35–42 in L1 larvae. Plotted are cumulative distributions of mean mRNA fold changes observed upon loss of EBAX-1, comparing the effect on predicted targets (red, left), predicted conserved targets (red, right), or control mRNAs lacking sites to miR-35–42 (blue). Statistical significance of differences between fold-change distributions was evaluated using the Kolmogorov–Smirnov (K–S) test.

### miR-35–42 appear to undergo trimming as they age

In mammals and flies, miRNAs subject to TDMD are sometimes also subject to target-directed tailing and trimming (TDTT) (Ameres et al. 2010; de la Mata et al. 2015; Sheu-Gruttadauria et al. 2019). Co-occurrence of TDMD and TDTT is thought to result from the conformational changes in AGO and miRNA that occur upon pairing to extensively complementary target RNAs, which render AGO susceptible to ZSWIM8-mediated polyubiquitination and the miRNA accessible to nucleases and terminal nucleotidyltransferases (Sheu-Gruttadauria et al. 2019; Han et al. 2020; Shi et al. 2020). Although tailed and trimmed isoforms of ZSWIM8-sensitive miRNAs accumulate upon loss of ZSWIM8, perturbation of miRNA tailing does not detectably influence the rate of ZSWIM8-dependent degradation in mammalian and fly cells, indicating that TDMD and TDTT are coincident but independent phenomena in mammals and flies (Han et al. 2020; Shi et al. 2020; Kingston and Bartel 2021).

We wondered about the extent to which tailing and trimming accompanies the degradation of EBAX-1–sensitive miRNAs in worms. For the miR-35 family and any other EBAX-1–sensitive miRNAs for which the 3′-end sequences do not influence degradation, TDTT might not coincide with loss of EBAX-1, whereas for any miRNAs degraded through the canonical TDMD pathway, TDTT might coincide with EBAX-1 loss. To examine the effect of EBAX-1 on miRNA tailing and trimming, we quantified the occurrence of tailed and trimmed miRNA isoforms in our staged sRNA-seq datasets.

We first calibrated the sensitivity of this approach to errors introduced during library preparation and sequencing, examining miscalled tailing and trimming of two synthetic miRNA sequences (*Drosophila melanogaster* (dme)-miR-14 and *Xenopus tropicalis* (xtr)-miR-428) that we had spiked into our total RNA samples before library preparation. A maximum of 0.13% mono-adenylated and 0.06% mono-uridylated reads were observed for either synthetic RNA across all 28 staged sRNA libraries included in this analysis (Table S5). Moreover, no spike-matching reads with more than one non-templated adenosine or more than two non-templated uridines were observed in these libraries (Table S5). Likewise, only low levels of miscalled 3′ trimming were observed—at most 0.81% of reads trimmed by 1 nt and 0.14% by 2 nt (Table S5).

Although in early embryos loss of EBAX-1 did not substantially affect the fractional abundance of any trimmed or tailed miR-35 isoforms, differences began to arise in late embryos (Fig. 4A, Table S6). During wild-type embryogenesis, the proportion of full-length miR-35 declined only modestly from ∼84% in early embryo to ∼82% in late embryo and ∼78% in L1 larvae, whereas in the mutant, the proportion of full-length miR-35 declined from ∼82% in early embryos to ∼72% in late embryos and ∼62% in L1 larvae (Fig. 4A, Table S6). This difference was explained by an increase in the fraction of 3′-trimmed isoforms in mutant late embryos and larvae. Whereas in wild-type L1 larvae, ∼10% of miR-35 was trimmed by 1 nt and ∼2.5% was trimmed by 2 nt, in larvae lacking EBAX-1, ∼23% of miR-35 was trimmed by 1 nt and ∼6.6% was trimmed by 2 nt (Fig. 4A, Table S6). This accumulation of trimmed miR-35 isoforms was also detectable by northern blotting (Fig. 2D). The increased percentages of trimmed isoforms observed in the *ebax-1* mutant continued through the later larval stages (Fig. 4A, Table S6). For instance, in *ebax-1* mutant L3 larvae, ∼27% of miR-35 molecules were trimmed by 1 nt, ∼9% by 2 nt, and ∼3% by 3 nt, compared to ∼8%, ∼2%, and ∼0.5% in wild-type L3 larvae, respectively (Fig. 4A, Table S6). These differences between mutant and wild-type diminished in gravid adult worms, coincident with the onset of the bulk of miR-35–41 biogenesis (Fig. 4A).

**Fig. 4.**
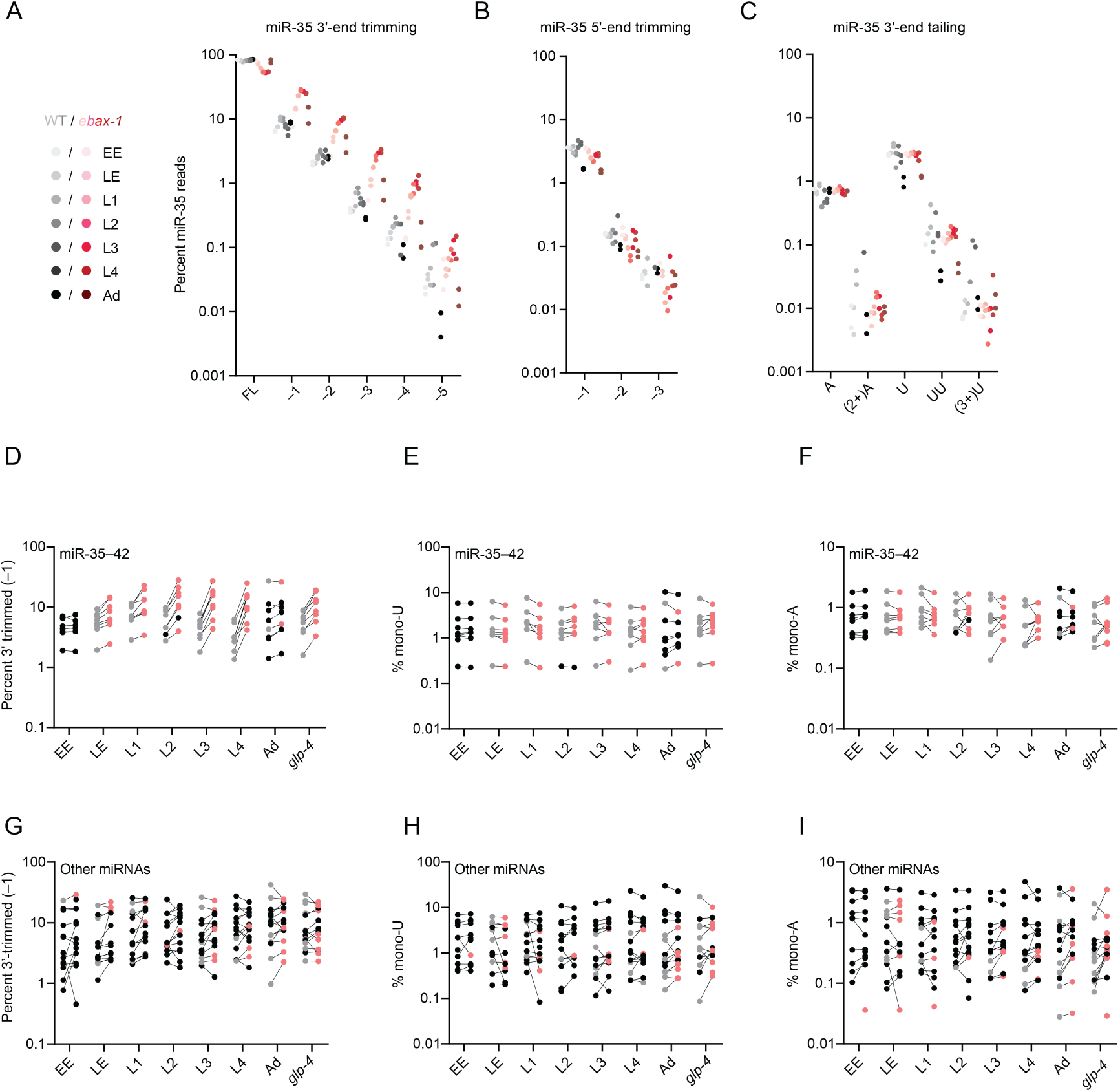
The miR-35–42 miRNAs that accumulate in *ebax-1* mutants undergo 3′ trimming. **(A–C)** The effect of EBAX-1 loss on the abundance of trimmed and tailed isoforms of miR-35 across worm development. Plotted are the percentages of miR-35 sRNA-seq reads that exactly matched full-length (FL) or 3’-trimmed isoforms **(A)**, 5’-trimmed isoforms **(B)**, or 3’ A-or U-tailed isoforms **(C)**, relative to the summed counts of all 14 of these isoforms. Points in gray show wild-type levels of each isoform; points in red indicate levels in *ebax-1* mutants, with darker colors denoting later developmental stages, as shown in the key on the left. Values for each of two biological replicates are plotted as separate points. **(D)** Trimming of the eight miR-35 family members across development. Plotted are the percentages of sRNA-seq reads to miR-35–42 3′-trimmed by 1 nt, calculated as in (A). For miRNAs that were EBAX-1–sensitive at a particular developmental stage, percent trimming is plotted for wild-type (gray points) or *ebax-1* mutant worms (red points) at that stage. Black points represent miRNAs that were classified as EBAX-sensitive but not at the indicated developmental stage. **(E)** Mono-uridylation of miR-35–42, plotted as in (D). **(F)** Mono-adenylation of miR-35–42, plotted as in (D). **(G)** Trimming of the 14 EBAX-1–sensitive miRNAs outside the miR-35 family across development, plotted as in (D). **(H)** Mono-uridylation of the 14 EBAX-1–sensitive miRNAs outside the miR-35 family, plotted as in (D). **(I)** Mono-adenylation of the 14 EBAX-1–sensitive miRNAs outside the miR-35 family, plotted as in (D).

In contrast to these differences in the fraction of trimmed 3′-ends, we detected no clear differences in the fractions of either trimmed 5′-ends or tailed 3′-ends in the *ebax-1* mutant (Fig. 4B, 4C). Thus, an increase in the fraction of trimmed 3′-ends was the predominant isoform-level effect of EBAX-1 loss on miR-35 across development. The other miR-35 family members behaved similarly as miR-35; upon loss of EBAX-1, late embryos and larvae had significantly higher percentages of 3′-end trimmed isoforms (Fig. 4D, Table S7) with little change in the prevalence of either mono-uridylated or mono-adenylated isoforms (Fig. 4E, 4F, Table S7).

With respect to EBAX-1–sensitive miRNAs outside the miR-35 family, a few had modestly increased trimming percentages upon loss of EBAX-1 (Fig. 4G). For example, miR-71 was EBAX-1 sensitive from the L2 to the L4 larval stage, during which time it underwent a ∼2-fold increase in the fraction of 3′ ends trimmed by 1 nt (Table S7). However, other miRNAs had reduced percentages of trimmed isoforms. Moreover, for those with increased trimming percentages, the increases often did not correlate with EBAX-1 sensitivity. Some miRNAs had strong EBAX-1 sensitivity without increased trimming. For example, miR-43 was elevated ∼7-fold in L4 larvae lacking EBAX-1 but underwent no significant change in the fraction of trimmed isoforms at L4 or any other stage (Table S7). Others had increased trimmed percentages during stages at which the miRNA was not EBAX-1 sensitive. For instance, miR-44/45 had an increase in the percentage of trimmed isoform at all stages examined (Table S7) yet was EBAX-1 sensitive in only the late embryo (Fig. 1A). Likewise, we found no relationship between miRNA tailing and EBAX-1 sensitivity, with a few miRNAs undergoing increased tailing upon loss of EBAX-1, a few others undergoing decreased tailing (Fig. 4H, 4I, Table S7).

In sum, for the EBAX-1–sensitive miRNAs outside the miR-35 family, we did not observe a convincing signal for increased tailing or trimming in the *ebax-1* mutant (Fig. 4G–I)—a signal that would have implicated involvement of the canonical TDMD pathway, in which the transcript that directs degradation pairs extensively to the miRNA 3′ region, promoting both TDMD and TDTT. Nonetheless, we cannot rule out the possibility that the canonical pathway functions in *C. elegans*, but no signal was observed because, in contrast to the tailing and trimming activities of other animals, the tailing and trimming activities of *C. elegans* do not recognize and modify 3′ ends of extensively paired miRNAs. With respect to the miR-35 family, we suspect that the clear signal for increased trimming in the ebax-1 mutant (Fig. 4D) was not a consequence of extensive pairing to the miRNA to the 3′ region, because such pairing is not required for degradation of the miR-35 family (Donnelly et al. 2022). Instead, we suggest that the increased percentages of trimmed isoforms observed in the *ebax-1* mutant are a consequence of the increased lifespans of these miRNAs in the absence of EBAX-1. The increased lifespans would presumably increase the average age of the miR-35 family members in the late-embryo and larval stages, and because the fraction of tailed or trimmed isoforms can increase as *C. elegans* miRNA molecules age (Vieux et al. 2021), the increased lifespans could thereby lead to an increased fraction of trimmed isoforms. A substantial influence of lifespan on the fraction of older molecules requires a strong shutoff of new miRNA production—dynamics that might uniquely cause substantial change in the age of miR-35 family members but not in that of other EBAX-1–sensitive miRNAs.

### The miRNA 3′ region contributes to *ebax-1*-mediated destabilization of miR-43

The known examples of TDMD in mammals and flies each require that the miRNA 3′ region pairs extensively to a trigger RNA (Han and Mendell 2023; Buhagiar and Kleaveland 2024), In contrast, for miR-35 in *C. elegans*, the sequence of its 3′ region is dispensable for its clearance at the embryo-to-L1 transition (Donnelly et al. 2022). This observation implies that if EBAX-1 is destabilizing miR-35 through a target-directed mechanism, it is through a nonconventional destabilization mechanism that does not involve pairing of the miRNA 3′ region. This possibility raises the question of whether this nonconventional, 3′-independent mode of TDMD might generalize to all other EBAX-1–sensitive miRNAs. Alternatively, EBAX-1 might have retained the ability to mediate conventional, 3′-dependent TDMD in *C. elegans*, and the 3′-independent mode might apply to only a subset of the EBAX-1–sensitive miRNAs. To begin to address this question, we examined whether the EBAX-1–dependent degradation of miR-43 requires the sequence of its 3′ region, in an experiment modeled after that which uncovered the 3′ independence of miR-35 destabilization (Donnelly et al. 2022).

miR-43 is a member of the miR-2 family, which in *C. elegans* also includes miR-2, miR-250, and miR-797 (Fig. 5A). Like the miR-35 family, the miR-2 family has divergent 3′ sequences (Fig. 5A), yet with the exception of miR-2, each miRNA in this family was sensitive to EBAX-1 loss (Fig. 1A and B, 5B). Of the three EBAX-1–sensitive family members, miR-43 was the most highly expressed (Table S1) and the most sensitive to EBAX-1 loss—accumulating to 7.2-fold above wild-type levels in L4 animals (Fig. 5B, Fig. 1A, Table S1). Moreover, miR-43 is processed from the same primary transcript as miR-42 and miR-44, which provided additional possibilities for monitoring transcriptional effects (Fig. 5C).

**Fig. 5.**
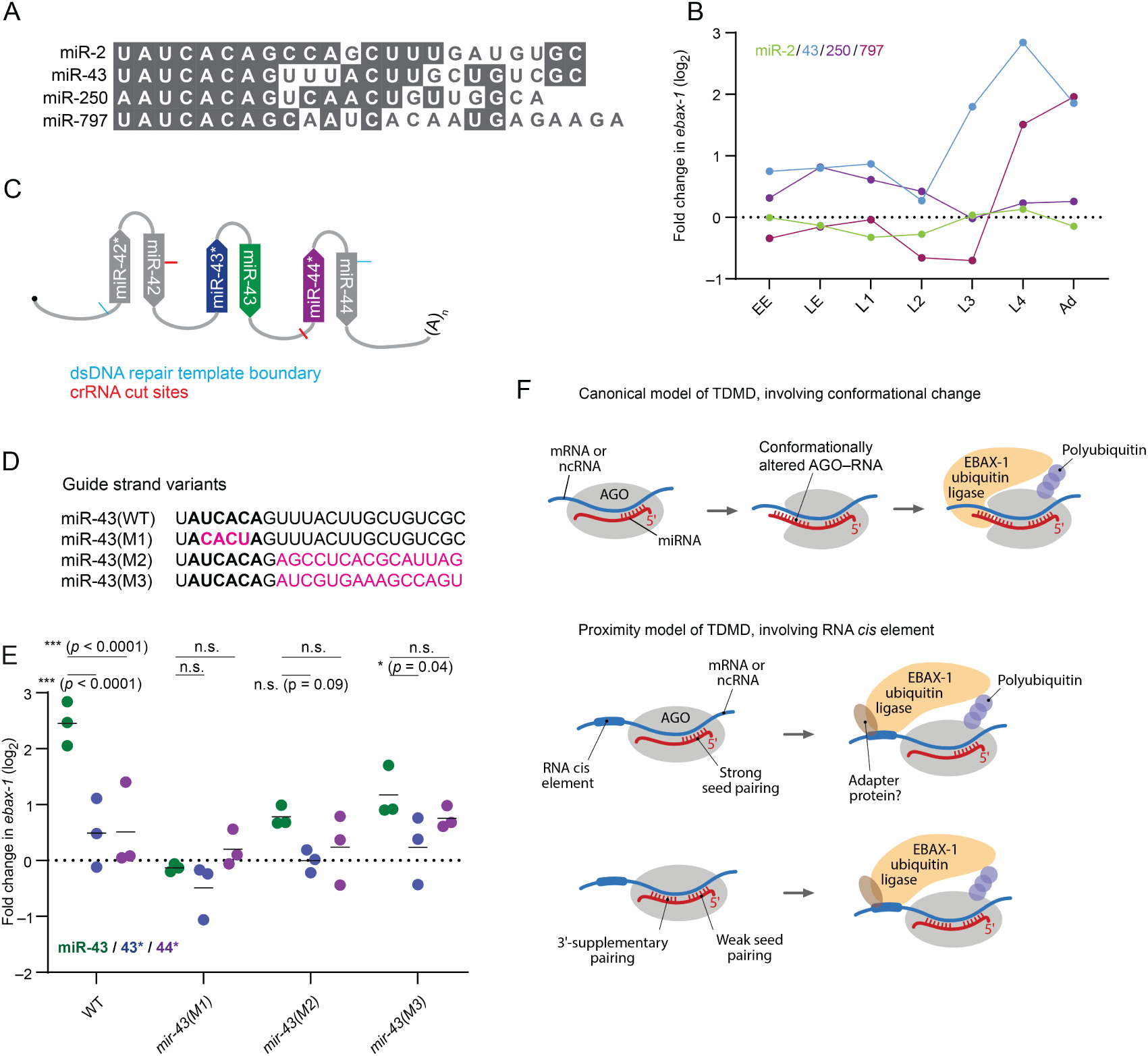
The miR-43 3′ region contributes to EBAX sensitivity. **(A)** Sequence alignment of the four members of the *C. elegans* miR-2 family. **(B)** EBAX-1 sensitivity of the miR-2 family members. **(C)** Schematic of the miR-42–44 cluster, indicating Cas9 cut sites (red lines) and repair template boundaries (blue lines) used for *in vivo* genome editing of this locus. **(D)** miR-43 variants generated through genome editing. Seed nucleotides are in bold, substitutions are in magenta. **(E)** The effect of EBAX-1 loss on the abundance of the miR-43 variants shown in (D) at the L4 stage. Points show fold change upon EBAX-1 loss in each miR-43 guide-strand variant (green), each miR-43 passenger-strand variant (blue), or each miR-44 passenger strand (purple) in wild-type worms or worms carrying the indicated *mir-43* mutations. Plotted are values for each replicate, with horizontal lines indicating mean values of replicates. To assess statistical significance, we performed a two-way ANOVA followed by Dunnett’s multiple comparisons test. n.s., not significant. *, p < 0.05. ***, p < 0.0001. For one of three replicates in worms without miR-43 mutations, the fold change in miR-43, miR-43*, and miR-44* represents the mean of L4-stage data in the sRNA-seq experiment shown in Fig. 1 and Table S1. **(F)** Molecular models for EBAX-1–mediated miRNA destabilization. In the canonical model of TDMD (top), which is consistent with observations in mammalian and insect cells, extensive pairing between the miRNA 3′ region and a trigger RNA (either an mRNA or ncRNA) causes a conformational change in AGO, leading to the recruitment of EBAX-1 and subsequent polyubiquitination of AGO. In an alternative model (bottom), which is consistent with observations for miR-35–42 in *C. elegans*, EBAX-1 is recruited to the trigger RNA by an RNA cis-acting element, either directly or with the help of an adapter protein. In this alternative model, either strong seed pairing or weak seed pairing supplemented by pairing to the miRNA 3′ region recruits the trigger, thereby recruiting EBAX-1 to the vicinity of AGO-like proteins.

Using Cas9-faciliated homologous recombination, we mutated the endogenous *mir-43* locus to generated three strains bearing substitutions of either the miR-43 seed or its 3′ end (Fig. 5D). One mutant, designated *mir-43(M1)*, had an inverted miR-43 seed. A second mutant, *mir-43(M2)*, had positions 9–23 substituted with random sequence. A third mutant, *mir-43(M3)*, had positions 9–23 substituted with the 3′ region of miR-82. miR-82 is not EBAX-1 sensitive and its 3′ region was also used for the mutagenesis study of miR-35 (Donnelly et al. 2022). In each case, we also introduced compensatory mutations to the passenger strands to preserve the secondary structures of the pri- and pre-miRNAs (Fig. S4A). We further crossed these *mir-43* mutations into the *ebax-1(tm2321)* background, allowing us to test whether the seed or 3′ mutations affect EBAX-1–dependent miR-43 degradation.

We raised synchronous L4 populations of the three *mir-43* mutants, the three *mir-43;ebax-1* double mutants, as well as wild-type worms and *ebax-1* worms without any *mir-43* mutations, and assayed levels of small RNAs by sRNA-seq (Fig. 5E). Wild-type miR-43 was elevated 5.5-fold in L4 larvae lacking *ebax-1*, with only a 1.2-fold increase in its passenger strand (Fig. 5E, S4B, S4C), in line with our initial findings (Fig. 1A, 5B). This increase was also much greater than that observed for the passenger strand of miR-44, which like the miR-43 passenger strand was processed from the same primary transcript as miR-43 and was expected to be EBAX-1 insensitive, and thus provided another means to monitor potential changes in transcription. In contrast to miR-43, miR-43(M1), which had the substituted seed, was refractory to EBAX-1 (Fig. 5E, S4B, S4C), as would be expected for any process involving miRNA targeting after substituting the seed. The miR-43(M1) and miR-44 passenger strands also had only minor changes upon EBAX-1 loss in this strain (Fig. 5E, S4C, S4D), in agreement with a post-transcriptional explanation for the insensitivity of miR-43(M1) to EBAX-1 loss.

With respect to the mutants in the miR-43 3′ region, mir-43(M2) increased by 1.7-fold upon EBAX-1 loss, and miR-43(M3) increased by 2.4-fold. These increases were significantly less than the 5.5-fold elevation observed for wild-type miR-43 (Fig. 5E), indicating that the sequence of the miR-43 3′ region is critical for full EBAX-1 sensitivity. Thus, the seed-only degradation mechanism acting on miR-35 and presumably other members of the miR-35 family (Donnelly et al. 2022) does not act on all the EBAX-1–sensitive miRNAs of *C. elegans*.

Although much lower than the 5.5-fold increase observed for wild-type miR-43, the increase observed upon EBAX-1 loss for the miR-43(M3) variant was significantly greater than that observed for its passenger strand, and the increase observed for miR-43(M2) was also greater than that observed for its passenger strand, although the difference for miR-43(M2) was not statistically significant (Fig. 5E). This being said, the increases were each not significantly greater than that observed for the miR-44 passenger strand transcribed from the same locus (Fig. 5E), raising the possibility that transcriptional effects might have contributed to the increases observed for these two mutants of the miR-43 3′ region.

Overall, the substantially reduced EBAX-1 susceptibility of both 3′-region substitutions pointed to a contribution of the 3′ region of miR-43 for EBAX-1 sensitivity, reminiscent of canonical TDMD in flies and mammals. Whether or not the modest sensitivity that remained for these 3′-region substitutions might have also have been post-transcriptional will require further investigation.

## DISCUSSION

We present a stringently curated set of EBAX-1–sensitive miRNAs profiled across worm development. These results extend the confidently annotated EBAX-1–sensitive miRNAs to developmental stages beyond the gravid adult (Shi et al. 2020), providing statistical support for miRNAs suggested to be EBAX-1 sensitive in embryos (Shi et al. 2020; Donnelly et al. 2022) or mid-to-late larvae (Nahar et al. 2024) and identifying others not previously suspected to be EBAX-1 sensitive. In total, we annotate 22 miRNAs as EBAX-1 sensitive in at least one developmental stage, and show that this sensitivity can impact the abundance of predicted target mRNAs. Indeed, EBAX-1 sensitivity appears to underlie the instability of most short-lived miRNAs in the worm, as the EBAX-1–sensitive miRNAs reported here have some of the shortest half-lives of all worm miRNAs (Fig. S2D). Nonetheless, our set of 22 miRNAs might not be complete. Our study examined only worm populations grown under standard, well-fed conditions (with the exception of starved L1 larvae); additional miRNAs might be EBAX-1 sensitive under stresses and other conditions encountered in the wild.

The simplest explanation for the EBAX-1 sensitivity of these 22 miRNAs is that they are TDMD substrates. Thus, our study extends the implied scope of TDMD in *C. elegans* to 22 different miRNAs and to all major phases of *C. elegans* development. This being said, until the RNA transcripts that trigger the degradation of these miRNAs have been identified, the possibility that their EBAX-1 sensitivity might be through some other mechanism cannot be ruled out. Thus, for now, we classify these EBAX-1–sensitive miRNAs as probable, but yet-to-be-validated, TDMD substrates.

The current model of canonical TDMD proposes that pairing of the 3′ region causes a conformational change in the miRNA–AGO complex that favors ZSWIM8/EBAX-1 binding and AGO polyubiquitination (Han et al. 2020; Shi et al. 2020) (Fig. 5F). The EBAX-1–dependent yet 3′-region-independent degradation of miR-35 in *C. elegans* indicates that a mechanism that does not rely on 3′ pairing can also recruit EBAX-1 (Donnelly et al. 2022), and raises the question of whether in *C. elegans* TDMD always acts independently of the miRNA 3′-region.

This might be the case if, for example, EBAX-1 has lost the ability to recognize the 3′-pairing-dependent conformational change, and instead recognizes the trigger RNA (or a factor bound to the trigger RNA) (Donnelly et al. 2022). In this scenario, which we call the proximity model, the trigger would recruit both the miRNA and EBAX-1 and bring them into proximity with each other without a requirement for 3′-pairing and its associated conformational change (Fig. 5F). Our observations for mutants of miR-43 shows that its 3′-region is required for substantial EBAX-1 sensitivity, which shows that in *C. elegans*, EBAX-1 does not always act independently of the miRNA 3′-region. Nonetheless, these observations do not necessarily show that EBAX-1 recognizes a conformational change imparted by 3′ pairing; the proximity model is still possible. For example, the miR-43 TDMD trigger might operate through the proximity model by recruiting miR-43 through a site in which extensive 3′ pairing supplements weak seed pairing. In this scenario, disrupted 3′-pairing would dramatically reduce TDMD but perhaps not fully eliminate it, as seems to have occurred when substituting the 3′ region of miR-43 (Fig. 5E). The use of 3′-supplementary (or 3′-compensatory) pairing when directing miR-43 degradation would also impart specificity to miR-43 compared to its other family members, which might help explain why miR-2 was not EBAX-1 sensitive. Differentiating between the conformation and proximity models, and exploration of other questions regarding EBAX-1 recognition and function, will be facilitated with the identification of transcripts that trigger miRNA degradation in *C. elegans*.

## MATERIALS AND METHODS

### *C. elegans* strains, maintenance, and staging

Worms were cultured at 20°C on standard NGM agar media seeded with *E. coli* strain OP50 as described (Brenner, 1974), except for *glp-4(bn2)* and *glp-4(bn2); ebax-1(tm2321)* worms, which were maintained at either a permissive temperature of 15°C or a restrictive temperature of 25°C. The *ebax-1(ju699)* (strain CZ9912) was a gift from Yishi Jin. All double-mutant strains carrying the *ebax-1(tm2321)* allele were generated by mating *ebax-1(tm2321)* (strain CZ9907) hermaphrodites with males of the appropriate strain. Table S8 lists all worm strains used in this study.

Cultures were synchronized by treatment of mixed-stage worms with an alkaline bleach solution (0.6M NaOH and 12% v/v Clorox bleach). Embryos isolated immediately after bleaching were collected as early embryos, whereas embryos allowed to develop a further 6 h in M9 buffer (3 g/L KH_2_PO_4_, 6 g/L Na_2_HPO_4_, 0.5 g/L NaCl, and 1mM MgSO_4_•7H_2_O) with gentle agitation were collected as late embryos. Embryos allowed to further develop and hatch under these starvation conditions were collected as L1 larvae. Synchronous populations of older larvae were obtained by plating starved L1 onto NGM agar seeded with excess OP50 bacteria and allowing them to develop at 20°C until they reached the desired stage, which took approximately 16, 28, 42, and 68 h for L2, L3, L4, and gravid adults, respectively. *glp-4* and *glp-4; ebax-1* worms were hatched overnight at 15°C following bleaching, then synchronously raised at 25°C for approximately 48 h to reach adulthood.

### Sample collection and RNA extraction

Larvae and adult worms destined for RNA extraction were washed off of NGM agar plates into 15 mL tubes with M9 buffer. Worms were pelleted and washed twice with M9 buffer. Worms were then resuspended in M9 buffer and nutated a further ∼30 min at room temperature to purge bacteria from the gut. Worms were washed and centrifuged a final time, and the resulting pellet was flash-frozen in liquid nitrogen. Embryos and worms suspended in M9 buffer were pelleted and flash-frozen in liquid nitrogen.

Total RNA was extracted with TRI reagent (Life Technologies). Following the addition of TRI reagent to frozen worm pellets, the resulting slurry was flash-frozen and thawed twice more to disrupt the outer cuticle of the embryo or worm. Phase separation was achieved with 1-bromo-3-chloropropane (Sigma), and the resulting aqueous phase was precipitated in isopropanol. For embryo and larval samples, GlycoBlue coprecipitant (Invitrogen) was included as a carrier. Precipitated RNA was washed once in chilled 70% ethanol, resuspended in water, and quantified by Nanodrop spectrophotometry.

### Northern blotting

For each RNA sample, 3 µg total RNA was electrophoresed on a 20% polyacrylamide urea gel, transferred to a Hybond-NX membrane (GE Healthcare) in a semi-dry apparatus (Bio-Rad), and baked ∼1 h at 60°C in a solution of EDC (*N*-(3-dimethylaminopropyl)-*N*′-ethylcarbodiimide; Thermo Scientific) in 1-methylimidazole to crosslink RNA 5′ phosphates to the membrane. Membranes were pre-hybridized in Ultrahyb-oligo (Invitrogen), probed with radiolabeled LNA or DNA oligonucleotides (Table S9), and imaged with a Typhoon phosphorimager (Cytiva). Signal was quantified with ImageQuant TL software. A step-by-step protocol for small-RNA northern blots is available at http://bartellab.wi.mit.edu/protocols.html.

### sRNA-seq library preparation and analysis

Libraries were prepared from 2–10 µg total RNA essentially as described (Kleaveland, 2018), with the addition of spike-in RNA oligonucleotides in proportion to the total RNA content of each sample. A step-by-step protocol for sRNA-seq library preparation can be found at http://bartellab.wi.mit.edu/protocols.html. Libraries prepared for the *ebax-1* developmental time course were sequenced on the Illumina HiSeq platform with 50-nt single-end reads. Libraries prepared for the *mir-43* mutagenesis experiments were prepared with redesigned dual-index PCR primers (Table S9) and sequenced on an Illumina NovaSeq platform with 100 x 100 nt paired-end reads, with only read 1 being analyzed. Reads were trimmed of adaptor sequences using cutadapt (Martin, 2011) and fastx_trimmer (FastX toolkit: http://hannonlab.cshl.edu/fastx_toolkit/) then filtered for base quality using fastq_quality_filter with parameters -q 30 -p 100.

*C. elegans* miRNAs were counted by string-matching the first 19 nt of each read to a dictionary of curated miRNAs (Jan et al., 2011). Matching 19-nt prefixes rather than full-length sequences captured reads for miRNAs that had undergone tailing or minor trimming at their 3′ ends. miR-44 and miR-45, which are identical in sequence, were included in this dictionary as a single miRNA named miR-44. The passenger strands miR-1829b* and miR-1829c*, which are also identical in sequence, were not considered in our analyses. Resulting miRNA counts were merged into a table and passed to DESeq2 (Love et al. 2014) for differential expression analysis after removing counts corresponding to miRNA spike-in oligonucleotides. Normalized miRNA abundance was calculated from raw miRNA counts as counts per million miRNA-matching reads (CPM) plus one pseudocount, and only miRNAs with more than 5 CPM were carried forward for subsequent analyses.

*C. elegans* piRNAs (21U-RNAs) were counted by perfectly matching reads to a dictionary of 15,363 full-length piRNAs curated from ENSEMBL along with their genomic coordinates. These counts were then passed to DESeq2 for differential expression analysis. Only those piRNAs supported by at least one read in each wild-type and *ebax-1* sample were carried forward for plotting. Normalized piRNA abundance was calculated from raw piRNA counts as counts per million piRNA-matching reads (CPM), without adding a pseudocount.

For analyses of miRNA tailing and trimming, a dictionary of full-length miRNA sequences was modified to correspond to each altered miRNA species examined (for example, full-length miRNAs trimmed by 2 nt at the 3′ end). Perfect matches to these modified miRNA species (as well as unmodified full-length miRNAs) were tabulated. The relative abundance of each miRNA isoform is presented as the fraction of reads matching that isoform relative to the summed counts of all isoforms detected for that miRNA. Although the miRNA alterations analyzed were not an exhaustive catalog of all possible alterations, they did represent the most common modified miRNA species.

### Poly(A)+ RNA-seq library preparation and analysis

Poly(A)+ RNA-seq libraries were prepared from 3 µg total RNA from either late embryos or L1 larvae (for analysis of miR-35–42 target mRNAs) or from 10 µg total RNA from gravid adults (for analysis of protein-coding genes near dysregulated piRNAs on Chromosome IV) using the stranded NEXTFLEX Rapid Directional RNA-seq kit (Perkin Elmer). The embryonic and larval libraries represented two independent biological replicates of wild type and *ebax-1(tm2321)* samples, whereas the gravid-adult libraries represented one replicate. Libraries were all sequenced on the HiSeq platform with 50 nt single-end reads.

Reads were aligned to the *C. elegans* genome (WS245) using STAR v2.7.1a (Dobin et al., 2013) with parameters “ --outFilterMultimapNMax 1 --outFilterIntronMotifs RemoveNoncanonicalUnannotated --outSAMtype BAM SortedByCoordinate”. Uniquely mapping reads were assigned to genes using featureCounts (Liao et al., 2014) and parameters “-s 2 -F GTF -t CDS”, as well as *C. elegans* WS245 genome annotations downloaded from Wormbase (www.wormbase.org) and converted from gff3 format to gtf format with the gffread tool. Raw read counts were merged into a table and inputted to DESeq2 for differential expression analysis. Normalized mRNA abundance was calculated as counts per million mRNA-matching reads (CPM), without pseudocounting. A wild-type expression cutoff of 50 raw reads was used for plotting late-embryo and L1 mRNAs following differential expression analysis.

To analyze the effect of EBAX-1 loss on miR-35–42 target mRNA levels in late embryo and L1 (Fig. 3), sets of target and control non-target mRNAs were chosen as follows. All predicted mRNA targets of miR-35–42 were downloaded from TargetScanWorm and filtered for a wild-type expression threshold of 50 raw reads per mRNA in late embryo or L1 (separately for analysis of each time point). In addition to an analysis of all predicted miR-35–42 target mRNAs meeting the expression threshold, another analysis was performed for mRNAs categorized by TargetScanWorm as harboring “conserved” miR-35–42 sites. Sets of control nontarget mRNAs not listed in the miR-35–42 predictions were chosen (randomly and with replacement) for each target mRNA such that the length of their 3′ UTRs differed by no more than 10% (conserved) or 15% (predicted) from that of the predicted target mRNA, as defined by the longest 3′ UTR annotated for each mRNA in Jan et al. (2011). The distributions of fold changes in normalized mRNA abundance for target mRNAs and nontarget control mRNAs were then plotted as cumulative distribution functions using the Frequency Distribution analysis tool in GraphPad Prism 8 software, and analyzed for statistical significance by the Kolmogorov-Smirnov test.

### Cas9 facilitated substitutions

To construct the *mir-43(M1)*, *mir-43(M2)*, and *mir-43(M3)* lines, adult gonads were injected with in vitro-assembled RNPs containing recombinant Cas9 Nuclease V3 (IDT), alt-R tracrRNA (IDT), and alt-R crRNA (IDT) against the *mir-43* locus, as well as linear double-stranded DNA repair templates with homology arms of 100 bp. Our injection mixes also included a crRNA to the *dpy-10* locus and an ssDNA repair template to generate the dominant *dpy-10(cn64)* allele, in a “co-CRISPR” approach (Arribere et al. 2014) allowing for the selection of individuals that had taken up active Cas9 RNPs.

The injection mixes had 11.6 µM Cas9 nuclease V3, 11.6 µM alt-R tracrRNA, 4.8 µM of each of two alt-R crRNAs to the *mir-43* locus, 2.0 µM *dpy-10* crRNA, 1.4 µM *dpy-10(cn64)* ssDNA repair template, and 0.5 µM (86.8 ng/µL) dsDNA repair template for *mir-43* variants, in a total volume of 20 µL.

To prepare the injection mixes, crRNAs and tracrRNAs were first annealed by heating to 95°C for 5 min and cooling to room temperature. RNPs were then assembled by adding Cas9 nuclease V3 to this annealed duplex and incubating 5 min at room temperature. Repair constructs were then added. Each dsDNA repair template was purified by isopropanol precipitation and heat-denatured in parallel with the crRNA:tracrRNA mixture. Mixtures were centrifuged 5 min at 15,000*g* at room temperature in a benchtop microcentrifuge, and ∼18 µL was carefully pipetted from the top of the mixture to a new tube and loaded into needles pulled from capillary tubes for injection.

F1 worms displaying Dpy or Rol phenotypes, a function of their genotype at the *dpy-10* locus, were singled, and F1 worms were genotyped by Sanger sequencing of PCR amplicons whose boundaries spanned the region covered by the repair template. Homozygotes were then isolated from F2 or subsequent generations. Because *mir-43(M1)* mutants were identified in a background heterozygous for the allele imparting the Rol phenotype, F2 lines were screened for homozygosity at the miR-43 locus and for absence of the Rol phenotype. Because *mir-43(M2)* and *mir-43(M3)* lines were isolated in a *dpy-10* homozygous background, these two lines were backcrossed to wild-type N2 worms and then screened for homozygosity of the wild-type allele at the *dpy-10* locus as well as homozygosity of the appropriate mutant allele at the *mir-43* locus.

For genotyping, single worms were picked into PCR tubes containing 6 µL of a 5:1 mixture of Single-Worm Lysis Buffer (10 mM Tris pH 8.3, 50 mM KCl, 2.5 mM MgCl_2_, 0.45% IGEPAL CA-630, and 0.45% Tween-20) and Proteinase K (NEB, molecular biology grade) diluted tenfold in NEB storage buffer (20 mM Tris pH 7.4, 1mM CaCl_2_, and 50% glycerol). Alternatively, populations of F2 were occasionally genotyped by pipetting 30–50 µL of the same 5:1 lysis buffer:proteinase K mixture over a starved plate and recovering multiple F2 animals. In either case, the worm–buffer mixture was frozen and thawed twice on dry ice to disrupt the worm cuticle, then heated 1 h at 65°C to lyse, followed by 15 min at 95°C to inactivate proteinase K. To generate an amplicon for genotyping, 0.6 µL of worm lysate was used as template in a 15 µl PCR reaction, of which 5 µL was visualized on a 1% agarose gel and 10 µL processed for Sanger sequencing.

## ACKNOWLEDGMENTS

We thank members of the Bartel lab for helpful discussions, Y. Jin for some *C. elegans* strains, and S. Kennedy for providing access to his lab’s *C. elegans* microinjection space. D.P.B. is supported by NIH grant GM118135 and is an investigator of the Howard Hughes Medical Institute.

