## Supplemental Figures and Legends for "Widespread destabilization of *C. elegans* microRNAs by the E3 ubiquitin ligase EBAX-1"

### Figure S1

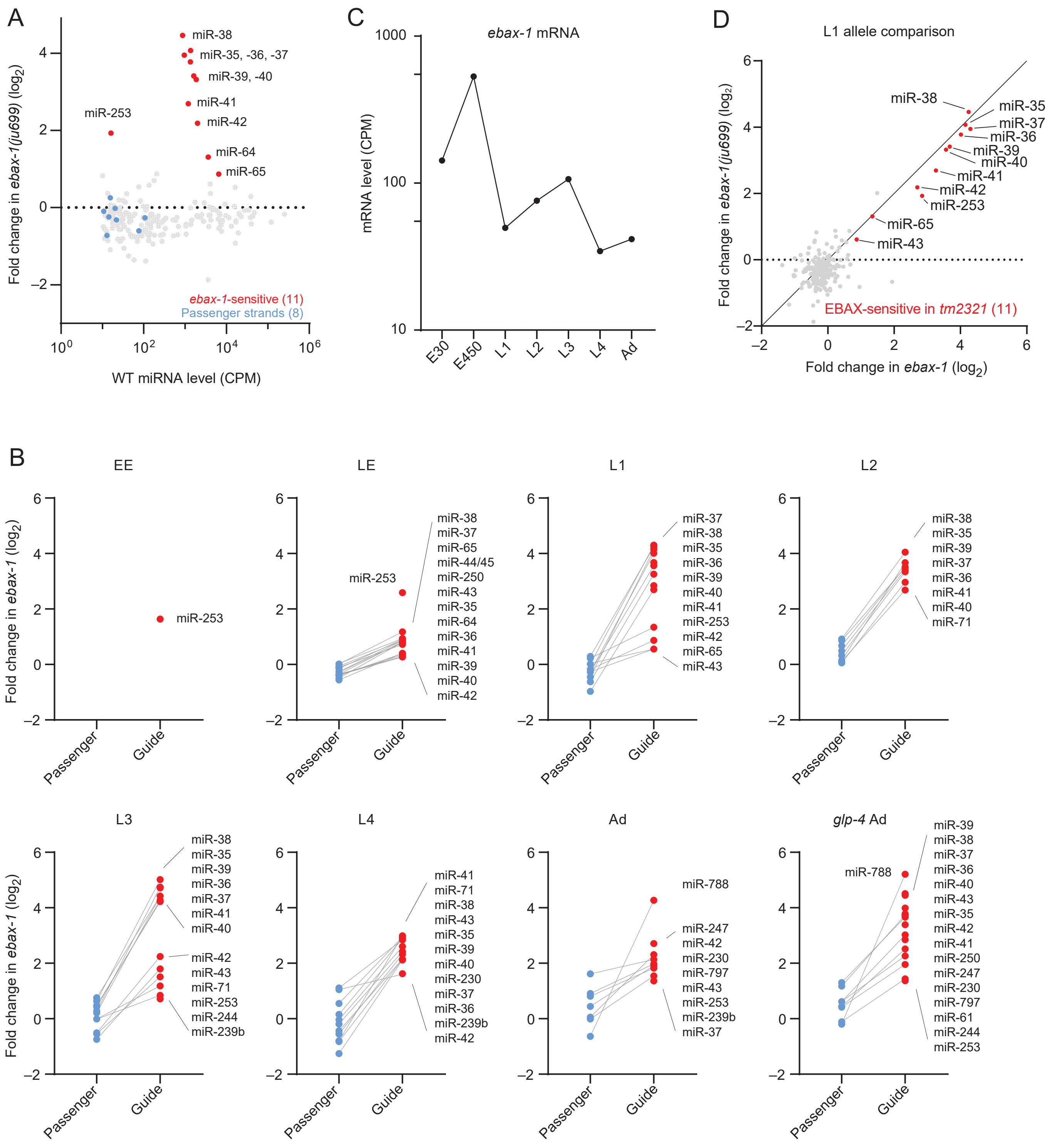
A
miR-38
miR-35, -36, -37
miR-39, -40
miR-41
miR-42
miR-253
miR-64
miR-65
ebax-1-sensitive (11)
Passenger strands (8)
Fold change in ebax-1(ju699) (log2)
WT miRNA level (CPM)
C
ebax-1 mRNA
mRNA level (CPM)
E30
E450
L1
L2
L3
L4
Ad
D
L1 allele comparison
Fold change in ebax-1(ju699) (log2)
Fold change in ebax-1 (log2)
EBAX-sensitive in tm2321 (11)
B
EE
LE
L1
L2
L3
L4
Ad
glp-4 Ad
Fold change in ebax-1 (log2)
Passenger
Guide
miR-788
miR-247
miR-230
miR-797
miR-239b
miR-250
miR-44/45
miR-71
miR-244
miR-61

### Figure S2

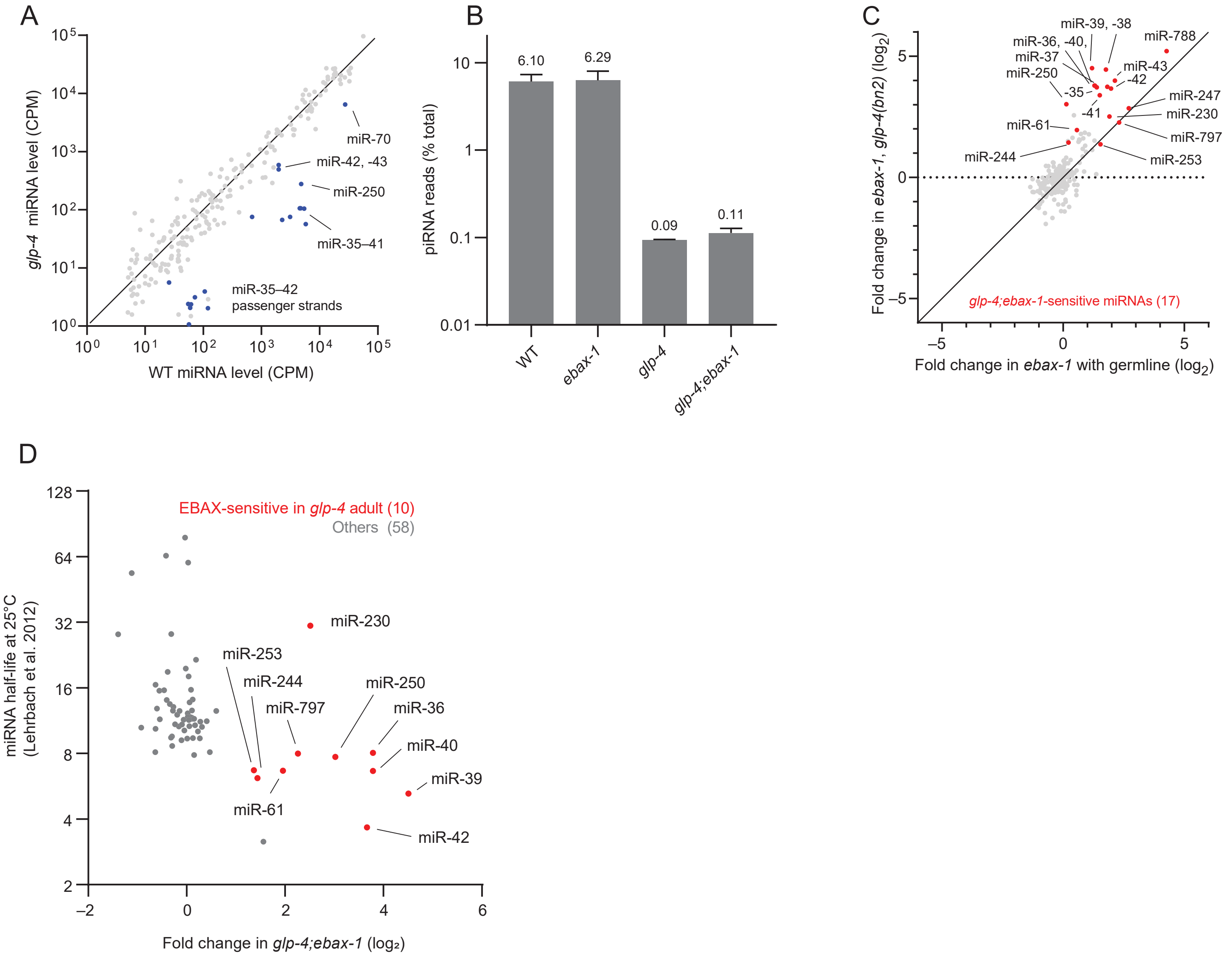

A
glp-4 miRNA level (CPM)
WT miRNA level (CPM)
miR-70
miR-42, -43
miR-250
miR-35–41
miR-35–42 passenger strands
B
piRNA reads (% total)
6.10
6.29
0.09
0.11
WT
ebax-1
glp-4
glp-4;ebax-1
C
Fold change in ebax-1, glp-4(bn2) (log2)
Fold change in ebax-1 with germline (log2)
miR-39, -38
miR-788
miR-36, -40,
miR-37
miR-250
miR-43
-42
-35
-41
miR-247
miR-230
miR-797
miR-61
miR-244
miR-253
glp-4;ebax-1-sensitive miRNAs (17)
D
EBAX-sensitive in glp-4 adult (10)
Others (58)
miRNA half-life at 25°C (Lehrbach et al. 2012)
Fold change in glp-4;ebax-1 (log₂)
miR-230
miR-253
miR-244
miR-797
miR-250
miR-36
miR-40
miR-39
miR-61
miR-42

Figure S3

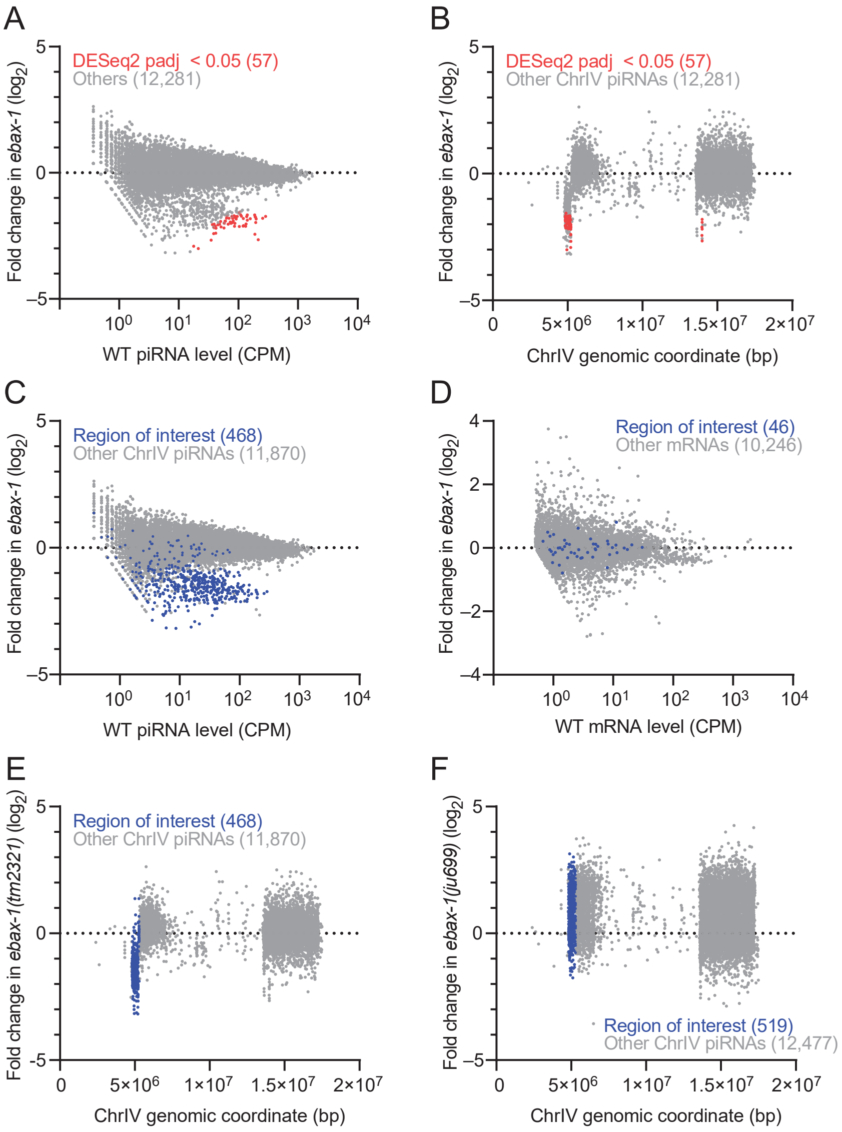

A
DESeq2 padj < 0.05 (57)
Others (12,281)
Fold change in ebax-1 (log2)
WT piRNA level (CPM)
B
DESeq2 padj < 0.05 (57)
Other ChrIV piRNAs (12,281)
Fold change in ebax-1 (log2)
ChrIV genomic coordinate (bp)
C
Region of interest (468)
Other ChrIV piRNAs (11,870)
Fold change in ebax-1 (log2)
WT piRNA level (CPM)
D
Region of interest (46)
Other mRNAs (10,246)
Fold change in ebax-1 (log2)
WT mRNA level (CPM)
E
Region of interest (468)
Other ChrIV piRNAs (11,870)
Fold change in ebax-1(tm2321) (log2)
ChrIV genomic coordinate (bp)
F
Region of interest (519)
Other ChrIV piRNAs (12,477)
Fold change in ebax-1(ju699) (log2)
ChrIV genomic coordinate (bp)

Figure S4

A

Passenger strand variants

miR-43(WT)* (gu)GACAUCAAGAAACUAGUGAUUAUG
miR-43(M1)* (gu)GACAUCAAGAAAUUAAGUGUUAUG
miR-43(M2)* (cu)AAUGAGUGAGCUCUAGUGAUUAUG
miR-43(M3)* (ac)UGGCCUUCAGAUCUAGUGAUUAUG

B

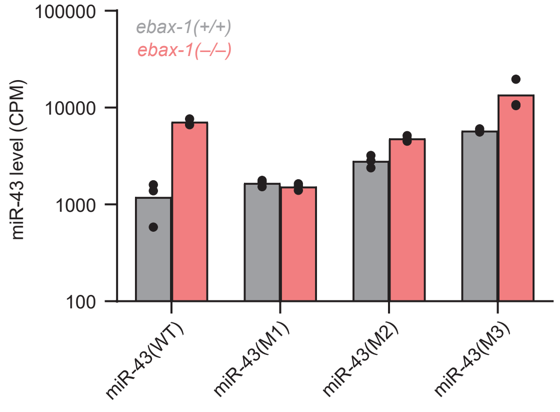


C

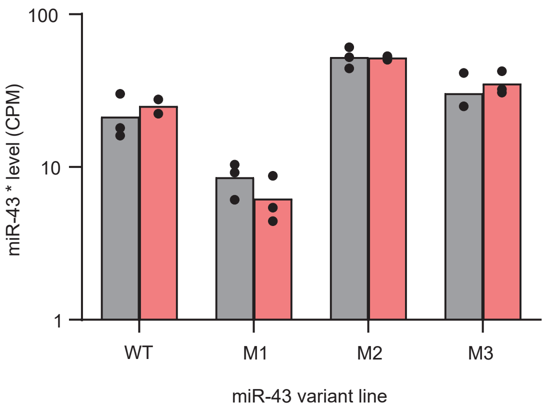


D

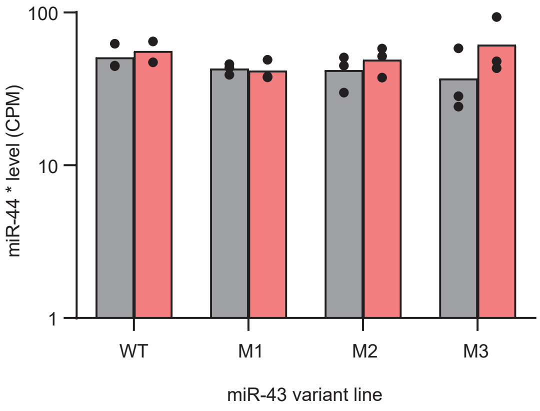

#### SUPPLEMENTAL FIGURE LEGENDS

**Fig. S1. sRNA-seq of *ebax-1* mutant worms across development. (A)** The effect of an independent null allele of *ebax-1* on miRNA levels in L1 larvae, plotted as in Fig. 1A. **(B)** Strand-asymmetric effect of EBAX-1 loss on miRNAs. Plotted are mean miRNA fold changes highlighted in Fig. 1, alongside the mean fold changes of the corresponding passenger strand(s). Passenger strands that did not meet our abundance threshold for reliable quantification are not shown. **(C)** The expression level of *ebax-1* mRNA over development. Raw mRNA counts were extracted from a poly(A)+ RNA-seq dataset (Boeck et al., 2016) and plotted in CPM. E30 denotes an early-embryo sample collected after 30 min of embryogenesis, whereas E450 denotes a late-embryo sample collected after 450 min of embryogenesis. mRNA levels in larval and adult stages are plotted as the mean of two biological replicates. **(D)** Comparison of two independent null alleles of *ebax-1* on miRNA levels in L1 larvae. Shown are mean fold changes of miRNAs in *ebax-1(ju699)* L1 larvae compared to wild-type larvae, plotted as a function of those changes in *ebax-1(tm2321)* L1 larvae.

**Fig. S2. Depletion of germline-enriched small RNAs in *glp-4* mutants. (A)** Comparison of miRNA levels in wild-type gravid adult worms raised at 20°C to those in *glp-4(bn2)* worms raised to adulthood at 25°C. Wild-type adult data correspond to those in Fig. 1. Points for miRNAs known to be depleted in germlineless worms (McEwen et al., 2016) are blue. **(B)** Comparison of piRNA levels in samples analyzed in (A). Shown is the mean percentage of piRNA-matching reads relative to all sRNA reads after adapter trimming and quality filtering. Error bars show the range between replicates. **(C)** The effect of germline loss on miRNA fold change in *ebax-1* mutants. The mean miRNA fold change due to EBAX-1 loss in germline-deficient adults raised at 25°C is plotted as a function of that observed in germline-intact adults raised at 20°C. Points corresponding to EBAX-sensitive miRNAs in the *glp-4* background are colored red. **(D)** The relationship between miRNA half-life and EBAX-1 sensitivity. *In vivo* miRNA half-lives measured in adult worms raised at 25°C (Lehrbach et al., 2012) are plotted as a function of the mean miRNA fold change observed upon EBAX-1 loss (as quantified in Fig. 1A) in *glp-4(bn2)* adults raised at 25°C. Points for EBAX-sensitive miRNAs are red.

**Fig. S3. A subset of piRNAs appears to be transcriptionally downregulated in *ebax-1(tm2321)*. (A)** Differences in piRNA levels observed in adult worms, as measured by sRNA-seq. Shown are mean fold changes of piRNA levels in *ebax-1(tm2321)* versus wild-type gravid adult worms across two biological replicates. piRNA levels are plotted as mean CPM. Results are shown for only piRNAs represented by at least one read in both wild-type and both *ebax-1* samples. Points for piRNAs meeting a DESeq2 adjusted *p*-value threshold of 0.05 are colored red. **(B)** Differences in piRNAs as a function of their location on Chromosome IV, which harbors nearly all piRNA genes in the worm. The mean fold change for each piRNA in (A) is plotted as a function of the starting coordinate of the corresponding piRNA gene, according to genomic coordinates downloaded from ENSEMBL Biomart. Points for piRNAs meeting a DESeq2 adjusted *p*-value threshold of 0.05 are colored red. **(C)** piRNA changes plotted as in (A), but indicating in blue the values for genes overlapping the interval ChrIV:4,800,000–5,300,000. **(D)** Differences in mRNA levels observed in adult worms, as measured by poly(A)+ RNA-seq. Plotted are fold changes of mRNAs in *ebax-1(tm2321)* gravid adult worms versus wild-type gravid adult worms, indicating in blue values for mRNAs overlapping the genomic interval ChrIV:4,800,000–5,300,000. mRNA levels are expressed as (CPM). Only values for mRNAs clearing a mean abundance threshold of 5 CPM in wild-type and *ebax-1* samples are plotted.

**(E)** Differences in piRNAs as a function of their genomic coordinate plotted as in (B), indicating in blue piRNAs overlapping the interval ChrIV:4,800,000–5,300,000. **(F)** Plotted as in (E), for an analogous experiment comparing the behavior of piRNAs in wild-type and *ebax-1(ju699)* gravid adults. The greater abundance of increases in piRNA levels is explained by asymmetric piRNA differences in the *ju699* and wild-type lines, in which an increase in the relative levels of many lowly expressed piRNAs is balanced by a decrease in the relative levels of some of the more highly expressed piRNAs.

**Fig. S4. The miR-43 3′ region influences its EBAX sensitivity. (A)** Passenger strands of miR-43 and its derivatives. Mutated positions are in magenta. Lowercase nucleotides in parentheses indicate pri-miRNA positions just upstream of the passenger strand that pair to the 3′-most positions of the guide strand. These positions were mutated in the M2 and M3 variants to preserve the secondary structure of *pri-miR-43*. **(B)** The effect of EBAX-1 loss on the abundance of the miR-43 variants at the L4 stage. Points plot the abundance of miR-43 variants from each of three biological replicates (two in the case of *ebax-1* larvae that were wild-type for miR-43). The height of each bar corresponds to the mean abundance of the indicated miR-43 variant in the presence (gray) or absence (red) of EBAX-1. miRNA abundance is plotted as CPM plus one pseudocount. **(C, D)** The effect of EBAX-1 loss on the passenger strands of miR-43 variants (C) and the passenger strand of miR-44 (D), in the *C. elegans* lines with different miR-43 variants, plotted as in (B).
