## Supplementary material for "Widespread destabilization of *C. elegans* microRNAs by the E3 ubiquitin ligase EBAX-1": Table Legends

**Table S1**. Changes in miRNAs observed upon loss of EBAX-1 during worm development, related to Figures 1, 2, S1, S2, and S3. Normalized miRNA counts, raw miRNA counts, and mean miRNA fold changes ($log_2$-transformed) are tabulated for wild-type and *ebax-1* mutant early and late embryo, L1, L2, L3, and L4 larvae, gravid adults, and germline-depleted adults. Samples designated strain CZ9907 correspond to *ebax-1(tm2321)* mutants, and strain CZ9912 corresponds to *ebax-1(ju699)*. Nomenclature such as N2_L1.1 versus N2_L1.2, and variants thereof, denotes biological replicates.

**Table S2.** Changes in piRNAs observed upon loss of EBAX-1 in gravid adult worms, related to Figure S3. Raw piRNA counts are tabulated for wild-type, *ebax-1(tm2321)*, or *ebax-1(ju699)* mutant samples. Samples designated strain CZ9907 correspond to *ebax-1(tm2321)* mutants, and strain CZ9912 corresponds to *ebax-1(ju699)*. Wild-type samples designated N2_A and N2_B were raised in parallel with *ebax-1(tm2321)* mutants, whereas wild-type samples N2_Ad_1 and N2_Ad_2 were raised in parallel with *ebax-1(ju699)* mutants in a separate experiment.

**Table S3.** Changes in mRNAs observed upon loss of EBAX-1 in gravid adult worms, related to Figure S3. Raw mRNA counts are tabulated for wild-type or *ebax-1(tm2321)* mutant samples, along with each mRNA's starting genomic coordinate and chromosome of origin.

**Table S4.** Changes in mRNAs observed upon loss of EBAX-1 in L1 larvae, related to Figure 3. The output of DESeq2 differential expression analysis of wild-type and *ebax-1(tm2321)* mutant miRNA counts is shown, and includes raw mRNA counts, normalized mRNA counts, and statistics such as adjusted *p*-value (padj).

**Table S5.** Trimming and tailing of synthetic oligonucleotides spiked into sRNA-seq libraries, related to Figure 4. For each oligonucleotide (dme-miR-14 and xtr-miR-428), tabulated are the mean, minimum, and maximum relative levels of each miRNA isoform across all sequencing libraries listed in Table S1. The isoforms included full-length (FL) molecules, molecules tailed at the 3′ end by one or multiple adenosines or uridines, and molecules trimmed at the 3′ or 5′ end by one or more nucleotides. Relative levels of each isoform are expressed as percentages of total reads matching all isoforms of a given oligonucleotide.

**Table S6.** Trimming and tailing of miR-35 across worm development, related to Figure 4. The relative levels of full-length (FL), trimmed, and tailed miR-35 isoforms are calculated as in Table S5. Relative isoform levels are tabulated for each individual sample represented in Table S1, with biological replicates grouped together. Raw counts corresponding to each miR-35 isoform are listed in a separate tab. Samples designated “ebax” correspond to *ebax-1(tm2321)* samples in Table S1.

**Table S7.** Trimming and tailing of the 22 EBAX-1 sensitive miRNAs across worm development, related to Figure 4. Relative levels of three miRNA isoforms – molecules trimmed by 1 nt at the 3′ end, mono-adenylated molecules, and mono-uridylated molecules – are tabulated in separate tabs and calculated as in Table S5 for each sRNA-seq library. Samples designated "CZ" are *ebax-1(tm2321)* mutant samples represented in Table S1.

**Table S8.** Worm strains used in this study.

**Table S9.** Oligonucleotides and CRISPR repair templates used in this study.
